# Sorting for CCR7-positivity enriches for a dendritic cell population with enhanced antigen-presenting capacity and anti-tumour potency

**DOI:** 10.1101/755488

**Authors:** Paul Burgoyne, Alan J Hayes, Rachel S Cooper, Michelle L Le Brocq, Christopher A.H. Hansell, John DM Campbell, Gerard J Graham

**Affiliations:** Chemokine Research Group, Institute of Infection, Immunity and Inflammation, University of Glasgow, 120 University Place, Glasgow, G12 8TA, UK; Scottish National Blood Transfusion Service, Jack Copland Centre, Research Avenue North, Edinburgh EH14 4BE, UK

**Keywords:** chemokines, dendritic cells, vaccines, tumours

## Abstract

Dendritic cell therapy has been a promising addition to the current armoury of therapeutic options in cancer for more than 20 years but has not yet achieved break-through success. To successfully initiate immunity, dendritic cells have to enter the lymph nodes. However, previous experience of therapeutic dendritic cell administration indicates that this is frequently an extremely inefficient process. The major regulator of dendritic cell migration to the lymph nodes is the chemokine receptor CCR7 and *in vitro* generated dendritic cells typically display heterogenous expression of this receptor. Here we demonstrate that positive-selection for the dendritic cell subpopulation expressing CCR7 enriches for cells with enhanced lymph node migration and antigen presentation competence as well as a chemokine expression profile indicative of improved interactions with T cells. In models of both subcutaneous and metastatic melanoma we demonstrate that the dendritic cells sorted for CCR7 expression trigger enhanced CD8 T-cell driven antitumour immune responses which correlate with reduced tumour burden and increased survival. Finally we demonstrate that this approach is directly translatable to human dendritic cell therapy using clinical-grade cell sorting.

**Synopsis:** Therapeutic dendritic cells drive anti-cancer immune responses but they migrate inefficiently to lymph nodes. We show that enriching for CCR7 expression yields a dendritic cellular product with enhanced ability to generate immune responses to localised and disseminated tumours.

## Introduction

Cellular therapy is an increasingly important treatment option in a range of clinical contexts including cancers and inflammatory/immune disorders(1-3). However, success is often limited by the mixed ability of the cells to home to, or be retained in, appropriate *in vivo* ‘therapeutic’ sites. In the context of dendritic cell (DC) therapy the extremely limited competence of therapeutically-administered DCs to home to lymph nodes (LNs) from the injection site (4-6) has contributed to a marked inefficiency of this therapeutic approach. Despite attempts to improve this, insufficient LN homing remains a frequent impairment to effective DC therapy. New approaches are therefore urgently needed to optimise such cell therapies.

Chemokines are central to in vivo tissue-homing of cells(7) and humans have approximately 45 chemokines many of which are expressed at select *in vivo* sites, or in specific inflamed contexts. Chemokines are broadly characterised and being inflammatory or homeostatic according to the in vivo contexts in which they function. The selective nature of chemokine expression ensures the attraction of cells bearing appropriate cognate chemokine receptors(8) to these destinations. One of the classic paradigms for chemokine receptor involvement in cellular homing focuses on CCR7(9), which is essential for cellular homing to, and positioning within, LNs(10). During DC maturation, upregulation of CCR7(9) ensures efficient LN migration to facilitate communication with T cells. Over-expression of CCR7 in DCs in murine models has been clearly linked with increased LN homing(11,12) and in human cancer patients, the extent of CCR7 expression on DCs correlates with the magnitude of the tumour infiltrating lymphocyte population(13). However, to date, attempts to increase LN homing competence of cells, either by manipulating CCR7 expression or by other means, have frequently been cumbersome and of limited applicability to the clinical context(11,12,14-16).

We have developed a novel approach to help address this issue(17). Whilst DC populations generated *in vitro* for therapeutic purposes invariably express heterogenous levels of CCR7, we have shown that selection of the subpopulation expressing CCR7 leads to a ‘fitter’ cellular product with enhanced LN homing. Typically such cell-selection would be carried out using antibodies but high quality antibodies are not available for the majority of chemokine receptors. We therefore use chemokines labelled with biotin or fluorophores, rather than antibodies, to select for cognate receptor-expressing cells(17,18). Specifically for sorting CCR7-expressing cells we use streptavidin-tetramerised biotinylated CCL19 (bCCL19).

Here we have used models of primary and metastatic cancer to show enhanced immune priming by CCR7-sorted DCs with improved outcomes in the tumour models. This study demonstrates the value of preselection of DCs for LN homing potential and highlights a potentially important novel development for therapeutic approaches to DC use in cancer vaccinology.

## Materials and methods

### Animals

C57BL/6 mice were from Charles River Laboratories, and OT-I mice were bred and maintained in house. All experiments were approved by the University of Glasgow Ethical Review Committee and performed under the auspices of a UK Home Office Licence.

### Isolation of mouse and human cells

Bone marrow-derived DCs (BMDCs) were generated as outlined in the Supplementary methods and as described previously(17). Murine T cells were isolated from harvested peripheral LNs of 8 week-old OT-I mice, by homogenising LNs through a cell strainer, and plated at 2×10^6^ cells/ml in mRPMI until use. Human buffy coats were obtained from SNBTS under appropriate sample governance and CD14+ monocyte-derived DCs (moDCs) generated as described previously(17) and in the Supplementary methods. Human T cells were maintained in culture by plating CD14-cells at 2×10^6^ cells/ml in hRPMI supplemented with 1µl/ml ConA (Sigma-Aldrich). On day 4, 20U/ml Xuri IL-2 (GE Healthcare) was added, and cultures maintained for up to 2 weeks. BMDCs and moDCs were sorted for CCR7 expression using bCCL19 as outlined in the Supplementary methods and as described previously(17).

### In vitro T cell stimulation

To assess DC stimulation, T cells were co-cultured with either CCR7-sorted or unsorted DCs, or 20U/ml IL-2 alone, at a ratio of 1:25 DCs:T cells in complete RPMI. Cultures were maintained for 7 days (mouse) or 14 days (human), with media replaced as necessary.

### Animal experiments – footpad migration

BMDCs were either CCR7-sorted or left sorted after undergoing the same labelling process as the sorted cells and labelled using either PKH67 dye (Sigma-Aldrich) or CellTracker Red (CMTPX) (Life Technologies) as per the manufacturer’s guidelines. CCR7-sorted or unsorted DCs were injected into the footpad of mice and after 48h the draining popliteal LNs and footpads were collected for enumeration of DCs in each tissue. To analyse cells present in the footpad, the tissue was digested using 2mg/ml Collagenase IV, 2mg/ml hyaluronidase and 0.1mg/ml DNase I (all Invitrogen) in RPMI for 20min at 37°C(19).

### Animal experiments – tumour models

To generate subcutaneous tumours, 8-week old male mice were injected on the right flank with 5×10^5^ B16.ova cells (B16 cells were obtained from ATCC). Tumours were allowed to grow until 12mm in diameter. To generate metastatic tumours, 8-week old male mice were injected intravenously with 5×10^5^ B16.ova cells and culled after 14 days. All tumour experiments had at least 5 animals per group.

### Statistical analyses

Data are presented as mean ± SEM. Statistical analyses were performed using GraphPad Prism 5 (GraphPad Software, La Jolla, CA, USA). Student’s t test and one-way ANOVA followed by Bonferroni’s multiple comparisons post-test were used. Tumour survival scores were assessed using Log-Rank (Mantel-Cox) test. Differences were considered statistically significant when p<0.05. p values <0.01 and <0.001 are also reported where appropriate.

***Additional Materials and Methods information is provided in the Supplementary information section.***

## Results

### Phenotypic characterisation of CCR7-sorted DCs

BMDCs display mixed expression of CCR7 (Figure 1Ai). CCR7+ cells were sorted from this mixed population using the bCCL19-sorting strategy(17) (Figure 1Aii). Henceforth, we refer to 3 cellular populations: i) unsorted DCs (‘mock’ sorted DCs); ii) sorted DCs (cells sorted for CCR7 expression) and iii) CCR7-DCs (the cells left from the unsorted population after removal of the CCR7+ cells). Flow cytometric analysis of F480 and CD11c staining demonstrated that, whilst CCR7-DCs distributed across 4 cellular populations, the sorted DCs were exclusively F480Lo/CD11cHi (Figure 1Aiii) and therefore represent a phenotypically-homogenous population of *in vitro*-derived DCs.

**Figure 1:**
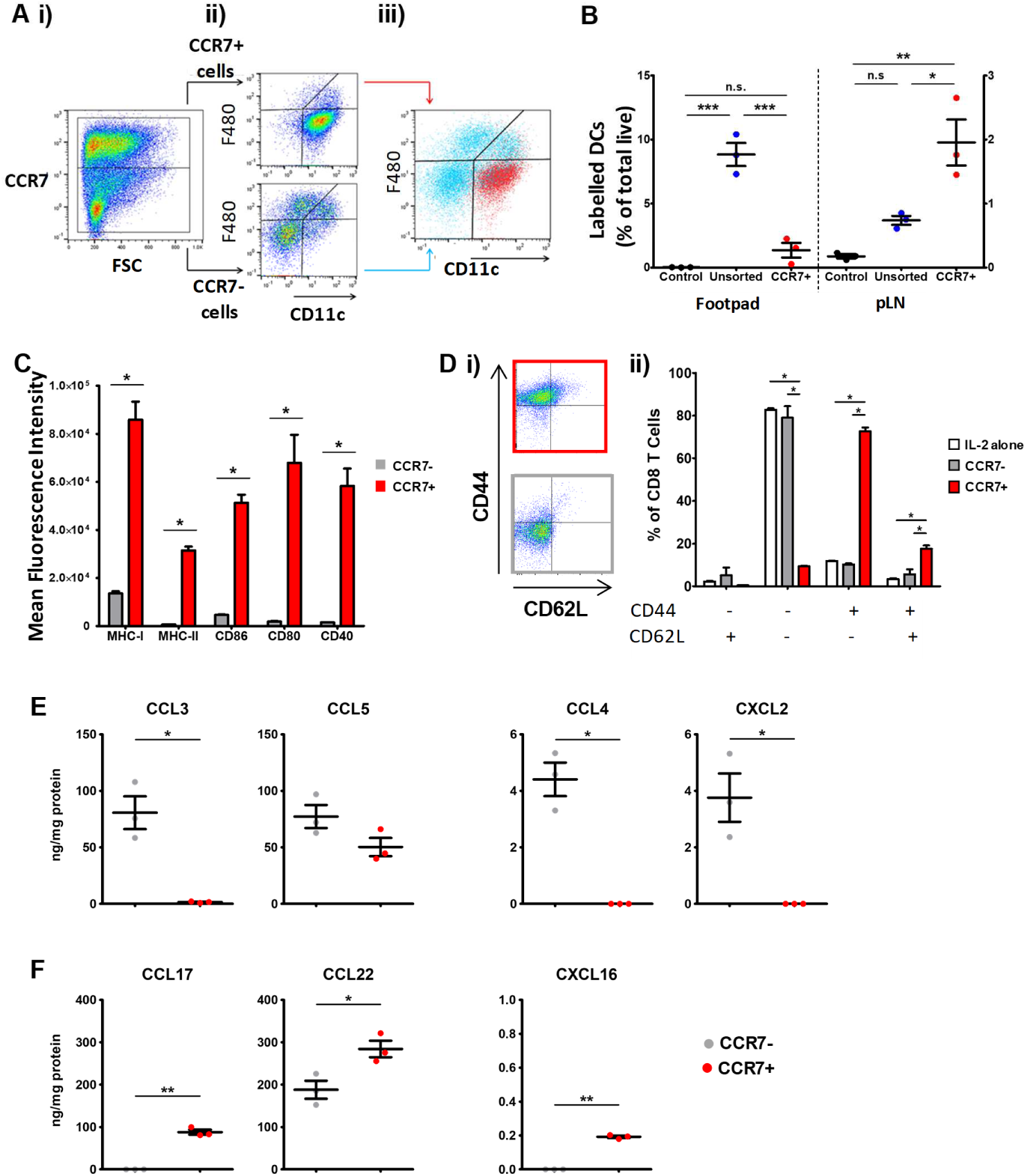
CCR7-sorted dendritic cells are enhanced for antigen presentation. A) i) BMDCs are heterogenous for CCR7 expression (FSC=Forward Scatter). ii) F480/CD11c flow cytometry profiles for CCR7+ve and -ve dendritic cells. iii) sorting for CCR7 expression isolates a discrete population of F480Lo/CD11cHi cells. B) numbers of the indicated dendritic cell populations remaining at the footpad injection site or migrating to the pLN. C) expression of markers of antigen-presenting competence in CCR7-ve (grey bars) and CCR7+ve (red bars) dendritic cells. D) flow cytometric analysis of CD44 and CD62L expression in T cells exposed to ova-pulsed CCR7+ve (upper panel) and CCR7-ve (lower panel) dendritic cells. E), F) expression of the indicated inflammatory (E) and T cell attracting (F) chemokines by CCR7-ve (grey dots) and CCR7+ve (red dots) dendritic cells. * p<0.05; * * p<0.01; * * * p<0.0001

In keeping with roles for CCR7 in cell migration to LNs(9), the sorted DC population migrated very efficiently from the footpad injection site to the draining popliteal LN (pLN) (Figure 1B). In contrast, the unsorted population, containing an equivalent number of CCR7+ cells, migrated poorly (Figure 1B). Next we examined the expression of conventional markers of DC maturity on the sorted cells. In contrast to CCR7-DCs, the sorted DCs expressed markedly higher levels of MHCI, MHCII, CD40, CD86 and CD80 (Figure 1C) indicating increased competency for antigen presentation and T cell stimulation. We used OTI CD8+ T cells to examine T-cell activating capacity of ova-pulsed sorted and CCR7-DCs. The sorted DCs preferentially induced a more mature CD44+CD62L-effector/memory CD8+ T cell phenotype whilst the CCR7-DCs supported less mature CD44-CD62L-CD8+ T cell populations (Figure 1Di and ii).

Luminex (Figure 1E) analysis revealed that, whilst CCR7-DCs expressed detectable levels of a number of inflammatory chemokines(20), these were either not expressed, or expressed at lower levels, by the sorted DCs. In contrast, the sorted DCs expressed elevated levels of chemokines binding to CCR4 (CCL17 and CCL22) and CXCR6 (CXCL16) (Figure 1F), which are involved in the attraction of effector/memory T cell populations, as well as regulatory T cells (21,22). These data therefore suggest that, in contrast to their CCR7-counterparts, the sorted DCs are more primed for development of a protective adaptive immune response through attraction of CD8+ and CD4+ T cells.

Together, these data demonstrate that sorting for CCR7+ DCs enriches for a population of cells with enhanced LN homing and T-cell activation competence.

### Sorted DCs display enhanced antitumour activity in vivo

We next tested the relative abilities of sorted, and unsorted, DCs to give rise to protective anti-tumour immune responses. As a model system we used the subcutaneous B16.ova melanoma model(23) which expresses ovalbumin as a tumour-specific neo-antigen.

Again, the number of unsorted cells injected into recipient mice was equivalent to the number required to give rise to the population of sorted DCs. In this model (Supplementary Figure 1A) ova-peptide pulsed DCs were injected into the footpad 1 day before subcutaneous tumour injection. Tumours developed for 17 days at which time all animals were culled and tissue harvested for analysis. Whilst the unsorted DCs did not alter tumour development, the sorted DCs had a significant effect on tumours with a 25% reduction in tumour size observable throughout the course of the experiment (Figure 2A). No differences were noted in time-to-initiation of tumour development. These data demonstrate that, in the context of an aggressively growing neoantigen-expressing tumour, sorted DCs are superior to unsorted DCs in mediating antitumour immune responses.

**Figure 2:**
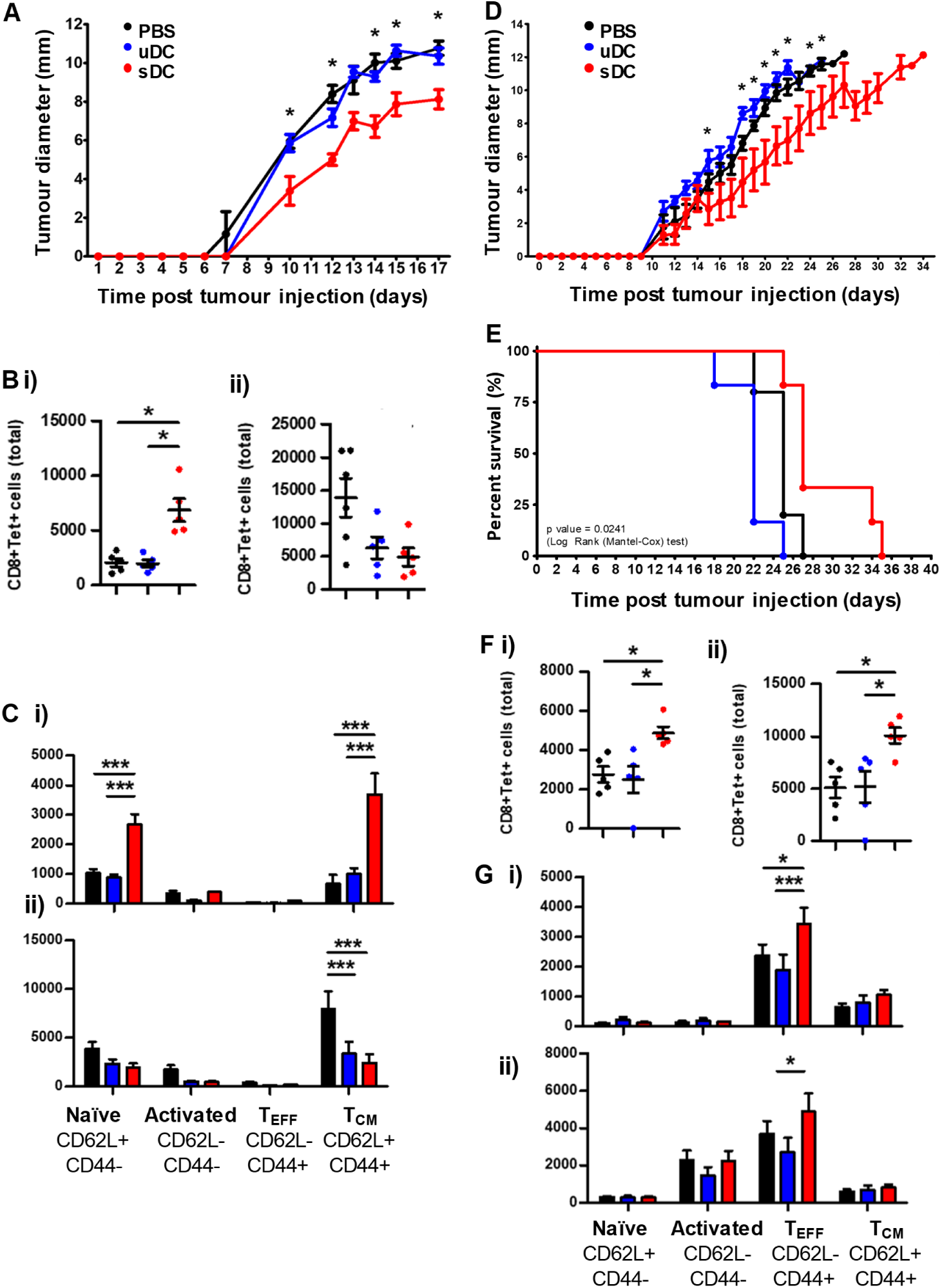
CCR7-sorted dendritic cells trigger enhanced antitumour immune responses. A) graph of subcutaneous tumour growth over the 17 days of the single dendritic cell injection model. B) numbers of ova-tetramer +ve CD8 T-cells in the i) draining popliteal and ii) non-draining inguinal lymph nodes. Black dots: PBS injected mice; blue dots: unsorted dendritic cell injected mice; red dots: CCR7+ve dendritic cell injected mice. C) phenotypic characterisation of T-cell populations in the i) draining popliteal and ii) non-draining iLNs. Black bars: PBS injected mice; blue bars: unsorted dendritic cell injected mice; red bars: CCR7+ve dendritic cell injected mice. D) graph of subcutaneous tumour growth over the 30 days of the double dendritic cell injection model. E) graph indicating mouse survival (time to cull) in a double dendritic cell injection model. Black line represents PBS injection; blue line represents unsorted dendritic cells and red line represents sorted CCR7+ve dendritic cells. F) numbers of ova-tetramer +ve CD8 T-cells in the i) draining popliteal and ii) non-draining iLNs. Black dots: PBS injected mice; blue dots: unsorted dendritic cell injected mice; red dots: CCR7+ve dendritic cell injected mice. G) phenotypic characterisation of T-cell populations in the i) draining popliteal and ii) non-draining iLNs. Black bars: PBS injected mice; blue bars: unsorted dendritic cell injected mice; red bars: CCR7+ve dendritic cell injected mice. * p<0.05; * * * p<0.0001

As shown in Figure 2Bi, and in keeping with the enhanced LN migration of DCs from the injection site, we found higher numbers of ova-specific CD8+ T cells in the footpad-draining pLNs of mice receiving sorted DCs compared to unsorted DCs. In contrast, we saw no difference in numbers of ova-specific CD8+ T cells in the non-draining inguinal LNs (iLN) (Figure 2Bii) thus the anti-ova response is specifically increased in the LNs draining the DC injection site. Notably, in the pLNs (Figure 2Ci) there was a specific increase in the numbers of ova-responsive naive and central memory CD8+ T cells. In contrast, there was a reduction in central memory T cell numbers in the iLNs of mice injected with both unsorted and sorted DCs (Figure 2Cii).

These data demonstrate that CCR7-sorted DCs have enhanced antitumour activity compared to unsorted cells associated with improved migration to draining LNs and induction of systemic central memory T cell responses.

### Double prophylactic administration of sorted DCs enhances antitumour responses

Next we examined whether 2 sorted DC administrations improved responses (Supplementary Figure 1B). Here, tumours were allowed to develop to a diameter necessitating animal cull allowing for a comparison of the relative ‘survival’ of mice receiving the sorted or unsorted DCs. We observed a 40-50% reduction in tumour development, as measured by tumour-diameter, in mice receiving the double prophylactic administration of sorted, compared to unsorted DCs or PBS control (Figure 2D). In addition (Figure 2E) this was associated with significantly increased survival of mice receiving sorted DCs compared to mice receiving either unsorted DCs or PBS control (p=0.0241). Analysis of ova-specifc CD8+ T cells indicated elevated numbers in the footpad-draining pLNs (Figure 2Fi) but increased numbers were also seen in the iLN (Figure 2Fii). Analysis of the specific CD8+ T cell phenotypes in these tissues (Figures 2Gi and Gii) revealed that they were predominantly of an effector phenotype.

Thus double prophylactic administration of sorted DCs is associated with a reduction in tumor development and increased ‘survival’. However, the antigen-specific T cell phenotype is now skewed in favour of an effector phenotype suggesting that repeated doses of CCR7+ DCs leads to the production of a greater number of effector T cells.

### Sorted DCs display enhanced antitumour activity in a model of pulmonary metastasis

The B16 model also lends itself to the development of pulmonary metastases (Supplementary Figure 2C) following intravenous injection(24). As shown (Figure 3A), the metastatic deposits that developed in mice receiving either PBS or sorted, or unsorted, DCs were histologically indistinguishable. However (Figure 3B), mice receiving sorted, in contrast to unsorted, DCs developed significantly fewer metastatic deposits in the lung. Indeed, we observed a 50% reduction in metastasis number in mice receiving sorted DCs compared to unsorted DCs or PBS administration. This was associated with increased numbers of ova-specific CD8+ T cells in the pLN (Figure 3C).

**Figure 3:**
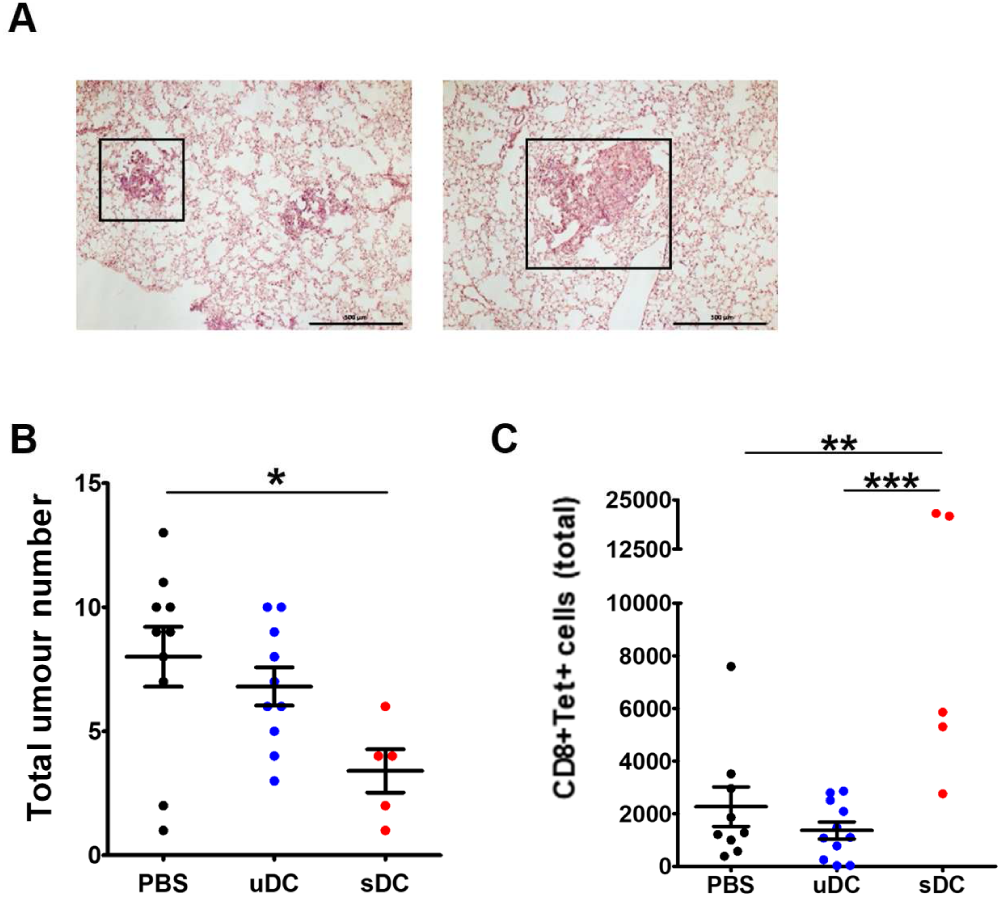
CCR7-sorted dendritic cells also enhance immune responses against metastatic tumours. A) exemplar histological analyses of pulmonary metastatic deposits from control injected (left panel) and CCR7-sorted dendritic cell injected (right panel) mice. B) measurement of metastatic burden in the lungs of mice as assessed in histological cross-sections of tumour bearing lungs. C) ova-tetramer positive CD8 T cell numbers in the popliteal lymph nodes draining the dendritic cell injection site. * p<0.05; * * p<0.01; * * * p<0.0001

These data demonstrate that the CCR7-sorted, compared to unsorted, DCs can induce an effective immune response against disseminated, metastatic cancers.

### This approach is translatable to human therapies

The use of cells in human therapy requires their production under the most stringent GMP conditions and the current methodology is suitable for translation to this standard using clinical-grade cell sorting. We have used a TYTO clinical-grade cell sorter (www.miltenyibiotec.com/upload/assets/IM0020121.PDF) and have adapted the bCCL19-sorting procedure for this machine. Using this clinical-grade sorter we were able to sort, with high efficiency and yield, large numbers of CCR7+ DCs from a mixed population of human moDCs (Figures 4Ai and 4Aii), the most commonly used *in vitro* source of therapeutic DCs. These sorted cells are viable (Figure 4Aiii) and, as with their mouse counterparts, express very high levels of molecules involved in antigen presentation and stimulation of T cells (Figure 4B). We next tested their ability to activate human T cells by incubating sorted and CCR7-DCs with autologous T cells for 10 days *in vitro* and then assessing T cell phenotypes using flow cytometry. In the context of EBV antigen presentation, there was limited change in the T cells but what was apparent was that the sorted DCs were capable of significantly enhancing the persistence of stem cell memory CD8+ T cells (Figure 4C).

**Figure 4:**
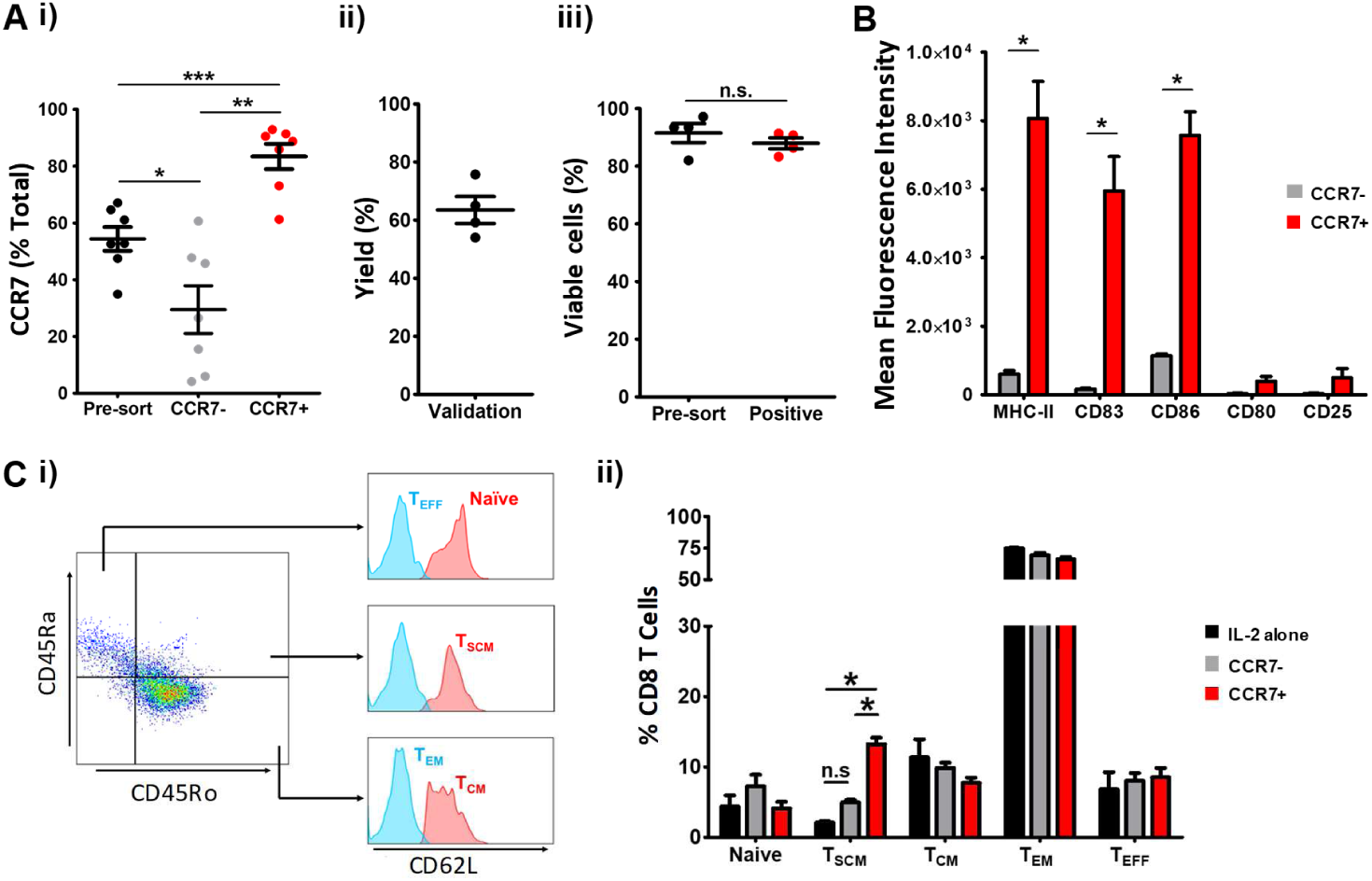
adapting the chemokine sorting protocol for clinical grade use. A) i) chemokine based sorting enriches moDCs from being approximately 55% CCR7+ve to in excess of 80% CCR7+ve. ii) yields of approximate 60% are routinely obtained following sorting. iii) sorted cells are approximately 90% viable. B) expression of markers of antigen presenting competence in CCR7-ve (grey bars) and CCR7+ve (red bars) moDCs. C) i) gating strategy showing flow cytometry gates for effector (Teff), stem cell memory (Tscm) and central memory (Tcm) T cells. ii) quantification of the different T cell populations showing a specific increase in Tscm cells in response to CCR7 +ve moDC. * p<0.05; * * p<0.01; * * * p<0.0001

Overall these data demonstrate that the experimental approach developed for sorting cells using CCR7 is of potential direct clinical applicability.

## Discussion

Here we show that CCR7-selected DCs have enhanced LN homing, antigen presentation and T cell stimulation capacity which correlate with increased protection in both the subcutaneous B16 melanoma model and the intravenous model of pulmonary metastasis. Importantly, our data show the CCR7-sorted cells are superior to a population of unselected cells containing the same number of CCR7+ cells. This indicates that non-CCR7 expressing cells actively antagonise DC function in vivo and further emphasises the value of the sorting procedure described for therapeutic DC therapy compared to the use of a population of DCs expressing heterogeneous levels of CCR7. *In vitro* data show CCR7+ DCs, compared to CCR7-cells, can uniquely prime antigen-specific naïve and memory T cells, both of which contribute to a strong anti-tumour response and are therefore highly desirable in a therapeutic context(25). In addition, we demonstrate enhanced expression of T cell chemoattractants by sorted DCs suggesting that they may be better equipped to mediate interactions with T cells upon arrival in the LN. Finally, we report that our methodology is entirely translatable to clinical grade cell sorting using a newly available GMP-compatible flow sorter.

