## Supplementary information for "Sorting for CCR7-positivity enriches for a dendritic cell population with enhanced antigen-presenting capacity and anti-tumour potency"

**Supplementary Information. Burgoyne et al.**

### Supplementary Materials and Methods

#### Generation of murine BMDCs

Bone marrow was harvested from the femur and tibiae of 8-12 week-old C57BL/6J mice and plated at 2x$10^6$ cells/ml in murine RPMI-1640 (ThermoFisher Scientific) (containing 10% FBS, 2mM L-glutamine, 100U/ml penicillin and 100µg/ml streptomycin (all Sigma-Aldrich)) (mRPMI) supplemented with 20ng/ml GM-CSF. Media was replaced on days 2 and 4. On day 6 of culture, media was replaced with mRPMI containing 5mg/ml ovalbumin (ova) (Sigma-Aldrich) for 4h, before addition of 100ng/ml LPS (Source Bioscience) and 50ng/ml TNFα (Miltenyi Biotec) overnight.

#### Generation of human monocyte-derived DCs.

PBMCs were separated into CD14+ and CD14- fractions using anti-CD14 microbeads (Miltenyi Biotec), as per the manufacturer's guidelines. Monocyte-derived DCs (MoDCs) were generated by plating CD14+ cells at 2x$10^6$ cells/ml in human RPMI-1640 (containing 5% AB serum (SNBTS), 2mM L-glutamine, 100U/ml penicillin and 100µg/ml streptomycin) (hRPMI) supplemented with 50ng/ml recombinant human GM-CSF (Miltenyi Biotec) and 15ng/ml recombinant IL-4 (Miltenyi Biotec). Media was replaced on days 2 and 4. On day 6 of culture, media was replaced with hRPMI containing EBV antigen (EBV consensus peptivator, Miltenyi Biotec; as per the manufacturer's guidelines) for 24h, before addition of 20µg/ml PolyI:C (Miltenyi Biotec) and 1µg/ml PGE2 (Sigma-Aldrich) for 48h.

#### Histology

Melanin-positive tumour lesions in the lungs were quantified by light microscopy prior to histology. To prepare lungs, tissue was fixed in 1% PFA overnight followed by immersion in 30% sucrose (ThermoFisher Scientific) in PBS overnight and then embedded in OCT (Tissue Tek). Tissues were cut to 8µm sections and stained using a standard haematoxylin and eosin protocol.

**Flow cytometry**

Single-cell suspensions were washed and stained in PBS containing 1% FCS/1% human albumin (for mouse/human cells, respectively) for flow cytometry analysis. Mouse cells were stained with the following fluorophore-conjugated monoclonal antibodies:
CD11c (X9-15), CD40 (3/23), CD44 (IM7), CD62L (MEL-14), CD8 (5H10-1), CD80 (16-10A1), CD86 (GL-1), F4/80 (BM8), MHC class I (AF6-88.5) (all BioLegend), and MHC class II (M5/114/15/2) (Miltenyi Biotec). Human cells used the following antibodies: CD11c (Bu15), CD25 (BC96), CD80 (2D10), CD83 (HB15e), CD86 (IT2.2) (all BioLegend), CD45Ra (T6D11), CD54Ro (UCHL1), CD8 (BW135/80), and MHC class II (L243) (all Miltenyi Biotec). Data were collected using the Miltenyi MACSQuant flow cytometer (Miltenyi Biotec, Germany) as mean fluorescent intensity and analysed using FlowJo (Tree Star, Ashland, OR, USA).

**Luminex**

BMDC lysates were prepared using NP40 Cell Lysis Buffer (ThermoFisher Scientific) and protein concentration determined by Pierce bicinchoninic acid (BCA) protein assay (ThermoFisher Scientific) as per the manufacturer’s guidelines. Lysates were then analysed using the Bio-Plex Pro™ Mouse Chemokine Assay (Bio-Rad) as per the manufacturer’s guidelines and the protein concentration of each analyte determined using the standard curves generated.

**Chemokine-sorting of DCs.**

Briefly, cells were incubated with 400ng/ml biotinylated CCL19 (bCCL19) complexed with phycoerythrin-conjugated streptavidin (SAPE) (BioLegend) for 30min at a concentration of $10^7$ cells/100µl in PBS with 2mM EDTA (Ambion). BMDCs were sorted by FACS using the BD Aria II (BD Biosciences), gating on PE-expressing cells. MoDCs were stained with a CD45 antibody (REA747) (Miltenyi Biotec) and sorted using the clinical-grade MACSQuant Tyto sorter (Miltenyi Biotec), gating on CD45+ and PE-expressing cells.

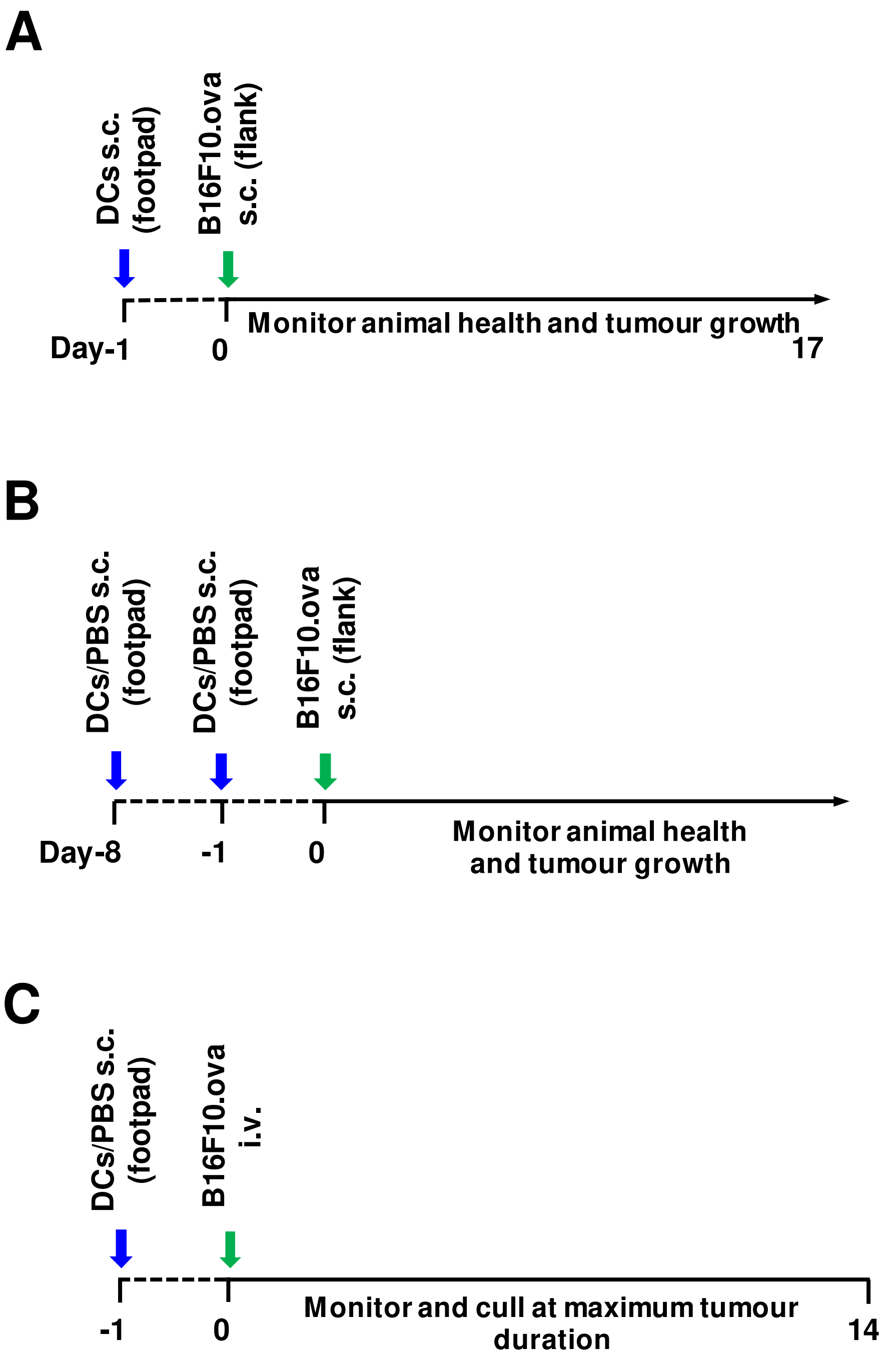


**Supplementary Figure 1.**

**Supplementary Figure 1: basic design of the in vivo experiments.**

A) the single dendritic cell injection model. Dendritic cells were injected into the footpad 24 hours prior to subcutaneous injection of B16F10-ova cells ($5x10^5$). Tumours were allowed to develop for 17 days at which point all mice were culled.

B) the double dendritic cell injection model. Dendritic cells were injected into the footpad 8 days prior, and 24 hours prior to subcutaneous injection of B16F10-ova cells ($5x10^5$). In this experiment tumours were allowed to develop to maximum permitted size to enable assessment of survival.

C) the metastatic melanoma model. Dendritic cells were injected into the footpad 24 hours prior to intravenous injection of B16F10-ova cells ($5x10^5$). Mice were culled at 14 days and lungs harvested for analysis.
